# Leucine suppresses glucagon secretion from pancreatic islets by directly modulating α-cell cAMP

**DOI:** 10.1101/2023.07.31.551113

**Authors:** Emily R. Knuth, Hannah R. Foster, Erli Jin, Matthew J. Merrins

## Abstract

**Objective:** Pancreatic islets are nutrient sensors that regulate organismal blood glucose homeostasis. Glucagon release from the pancreatic α-cell is important under fasted, fed, and hypoglycemic conditions, yet metabolic regulation of α-cells remains poorly understood. Here, we identified a previously unexplored role for physiological levels of leucine, which is classically regarded as a β-cell fuel, in the intrinsic regulation of α-cell glucagon release.

**Methods:** GcgCre^ERT^:CAMPER and GcgCre^ERT^:GCaMP6s mice were generated to perform dynamic, high-throughput functional measurements of α-cell cAMP and Ca^2+^ within the intact islet. Islet perifusion assays were used for simultaneous, time-resolved measurements of glucagon and insulin release from mouse and human islets. The effects of leucine were compared with glucose and the mitochondrial fuels 2-aminobicyclo(2,2,1)heptane-2-carboxylic acid (BCH, non-metabolized leucine analog that activates glutamate dehydrogenase), α-ketoisocaproate (KIC, leucine metabolite), and methyl-succinate (complex II fuel). CYN154806 (Sstr2 antagonist), diazoxide (K_ATP_ activator, which prevents Ca^2+^-dependent exocytosis from α, β, and δ-cells), and dispersed α-cells were used to inhibit islet paracrine signaling and identify α-cell intrinsic effects.

**Results:** Mimicking the effect of glucose, leucine strongly suppressed amino acid-stimulated glucagon secretion. Mechanistically, leucine dose-dependently reduced α-cell cAMP at physiological concentrations, with an IC_50_ of 57, 440, and 1162 μM at 2, 6, and 10 mM glucose, without affecting α-cell Ca^2+^. Leucine also reduced α-cell cAMP in islets treated with Sstr2 antagonist or diazoxide, as well as dispersed α-cells, indicating an α-cell intrinsic effect. The effect of leucine was matched by KIC and the glutamate dehydrogenase activator BCH, but not methyl-succinate, indicating a dependence on mitochondrial anaplerosis. Glucose, which stimulates anaplerosis via pyruvate carboxylase, had the same suppressive effect on α-cell cAMP but with lower potency. Similarly to mouse islets, leucine suppressed glucagon secretion from human islets under hypoglycemic conditions.

**Conclusions:** These findings highlight an important role for physiological levels of leucine in the metabolic regulation of α-cell cAMP and glucagon secretion. Leucine functions primarily through an α-cell intrinsic effect that is dependent on glutamate dehydrogenase, in addition to the well-established α-cell regulation by β/δ-cell paracrine signaling. Our results suggest that mitochondrial anaplerosis-cataplerosis facilitates the glucagonostatic effect of both leucine and glucose, which cooperatively suppress α-cell tone by reducing cAMP.

**Graphical Abstract:** 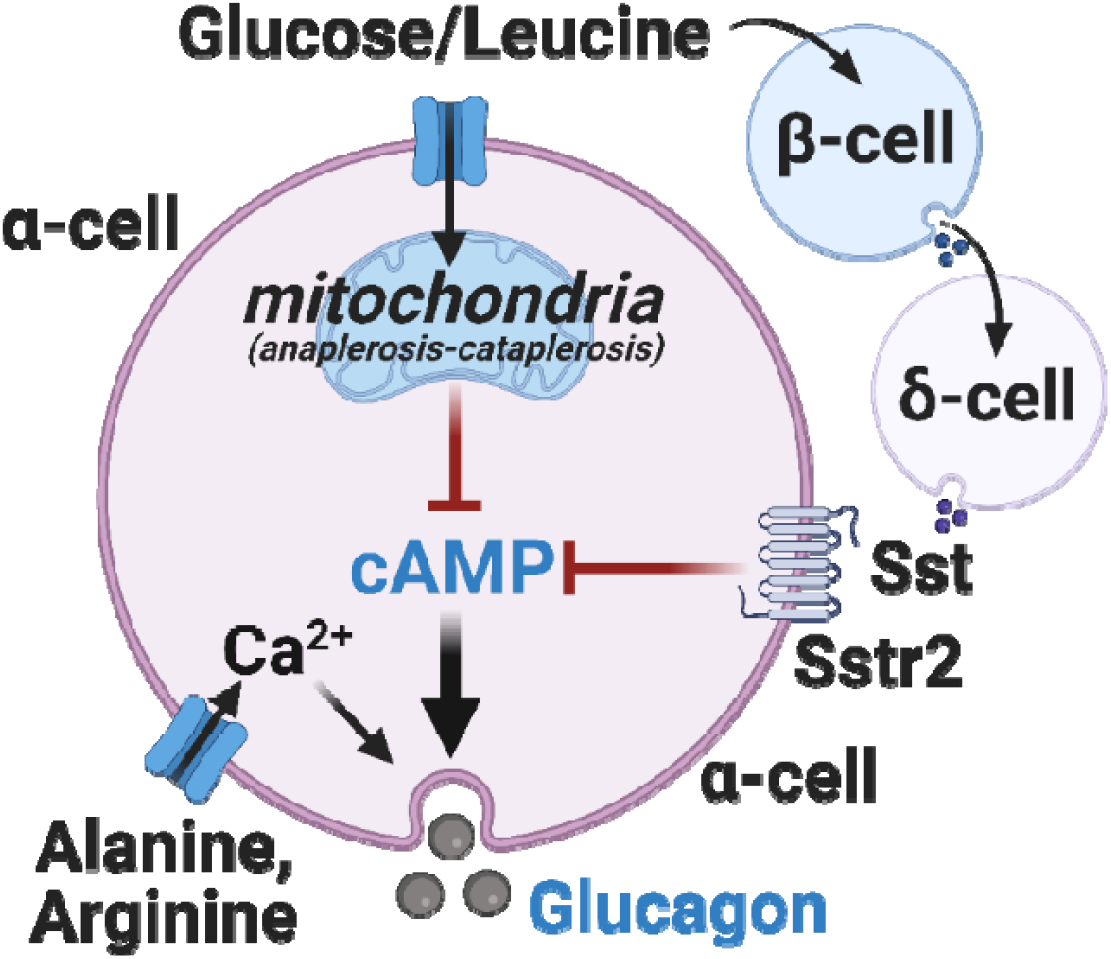

**Highlights:**

- Leucine inhibits glucagon secretion from mouse and human islets
- Leucine suppresses α-cell cAMP via both direct and paracrine effects
- Anaplerosis via glutamate dehydrogenase is sufficient to suppress α-cell cAMP
- Leucine suppresses α-cell cAMP and glucagon secretion more potently than glucose

## Introduction

Glucagon secretion from pancreatic islet α-cells plays a crucial role in maintaining blood glucose under hypoglycemic conditions [1–3], and supports insulin secretion in the fed state via glucagon signaling to neighboring β-cells [4–8], functions that are dysregulated in diabetes [9– 11]. Glucagon is released through Ca^2+^-dependent exocytosis that is regulated by the electrical activity of the α-cell [12–16]. Although Ca^2+^ itself does not have strong control over the magnitude of glucagon secretion [17–20], glucagon secretion is highly correlated with the level of α-cell cAMP, which acts by directly stimulating the exocytotic machinery and by recruiting additional glucagon granules into the readily releasable pool [15,21–23]. While these studies establish the respective roles of α-cell Ca^2+^ and cAMP in regulating the timing and magnitude of glucagon secretion, nutrient regulation of α-cell activity remains poorly understood.

Glucose is an inhibitor of glucagon secretion that regulates α-cell activity through both paracrine and intrinsic mechanisms. At elevated glucose concentrations, β-cell activation suppress glucagon secretion by releasing several paracrine factors that include insulin, GABA, zinc, and serotonin [21,24–27]. Insulin, despite reducing α-cell cAMP immunofluorescence [21], was later found to reduce glucagon secretion independently of cAMP [28,29], while UCN3 that is co-released with insulin suppresses α-cell cAMP by stimulating somatostatin release from δ-cells [21,30,31]. Antagonism of somatostatin receptor 2 (Sstr2), the primary α-cell somatostatin receptor in both mouse and human islets, increases glucagon secretion at both low and high glucose [21,28,32], establishing the importance of both tonic and glucose-stimulated paracrine inhibition by δ-cells. However, glucagon is maximally inhibited by glucose at concentrations that do not stimulate insulin release, including in somatostatin knockout mice [14,33,34], highlighting the importance of α-cell intrinsic regulation by glucose.

Intrinsic α-cell regulation by metabolic fuels remains an active but incompletely understood area of islet biology. Substantial research has focused on the ability of glycolysis, and glucokinase in particular, to suppress glucagon secretion [12,34–37]. Part of the glucagonostatic effect of glucose may be due to direct regulation of the K_ATP_ channel and α-cell electrical activity [12,14,33], which occurs partly or wholly via a K_ATP_-associated glycolytic metabolon [38]. In addition, H_2_O_2_ production in the mitochondria reduces P/Q-type Ca^2+^ channel activity and exocytosis [39]. Recent evidence also suggests an important role for mitochondrial anaplerosis (i.e., the net filling of the TCA cycle), which drives citrate cataplerosis (i.e., egress from the TCA cycle), producing cytosolic malonyl-CoA that switches off β-oxidation via inhibition of CPT1-dependent fatty acid transport [40,41]. Finally, via an unclear mechanism, glucose suppresses α-cell cAMP and glucagon secretion independently of Ca^2+^ regulation [28]. Collectively, these studies indicate that intermediary metabolism plays a crucial role in limiting glucagon secretion, although the majority have been conducted with glucose as the only substrate.

In addition to glucose, α-cells are sensitive to most amino acids [42]. For example, glucagon release is potently stimulated by the electrogenic fuels arginine (which is cationic) and alanine (which is cotransported with Na^+^) that depolarize the α-cell plasma membrane and stimulate Ca^2+^ influx [7,13,43]. Glutamine is not a potent secretagogue but is important for α-cell proliferation [44], while glutamate and glycine stimulate glucagon secretion through α-cell GPCRs [45,46]. One amino acid that has not received significant attention is leucine, which is primarily considered a β-cell fuel. Leucine is the only amino acid that is sufficient to stimulate insulin secretion, and like glucose, is both an oxidative and an anaplerotic mitochondrial fuel. While glucose and leucine are both metabolized to acetyl-CoA to oxidatively fuel the TCA cycle, they stimulate anaplerosis via different mechanisms. Glucose carbons anaplerotically fuel the TCA cycle via pyruvate carboxylase, which adds one carbon to pyruvate to generate oxaloacetate. Leucine carbons are not directly anaplerotic (since leucine-derived acetyl-CoA is lost in the TCA cycle as CO_2_), but leucine allosterically activates the anaplerotic enzyme glutamate dehydrogenase, adding carbon derived from glutamate to the TCA cycle as α-ketoglutarate. Gain-of-function mutations in glutamate dehydrogenase cause β-cell hypersecretion and protein-induced hypoglycemia [47]. However, it is not clear whether leucine regulates α-cell activity.

Here, we show that leucine suppresses glucagon secretion in both mouse and human islets. At physiological levels, leucine suppresses α-cell cAMP, without impacting α-cell Ca^2+^. Although leucine activation of β/δ cells contributes to its glucagonostatic effect, we find that leucine, similarly to glucose, functions independently of islet paracrine and juxtacrine signaling. Pharmacological activation of glutamate dehydrogenase is sufficient to mimic the effect of leucine, suggesting a central role for mitochondrial anaplerosis in suppressing glucagon secretion. Although glucose is also an anaplerotic fuel and suppresses cAMP, the much higher potency of leucine allows it to work more effectively under euglycemic and hypoglycemic conditions, when insulin secretion is low. Mechanistically, these data suggest a unifying function for mitochondrial anaplerosis in the regulation of α-cell activity through the intrinsic suppression of cAMP.

## Results

### Leucine suppresses glucagon secretion

Islet perifusion was used to simultaneously assess insulin and glucagon secretion from mouse islets stimulated with glucose (2 and 10 mM) and mixed amino acids provided at three times physiological concentration (LQRA, in mM: L, 1.5 leucine; Q, 1.8 glutamine; R, 0.6 arginine; A, 6.3 alanine) [7]. At low glucose, LQRA induced a large, biphasic rise in glucagon secretion that was further augmented by 10 nM glucose-dependent insulinotropic peptide (GIP) (Fig. 1A), consistent with the glucagonotropic effects of arginine/alanine and stimulation of the α-cell GIP receptor [7]. LQRA and GIP-stimulated glucagon release was dramatically reduced by elevated glucose (Fig. 1A). LQRA was sufficient to stimulate insulin secretion at low glucose, and amplified insulin secretion at high glucose (Fig. 1B). Since the glucagonostatic effect of glucose alone is small when compared with glucose suppression of amino acid-stimulated glucagon secretion, we used this latter approach to isolate the impact of leucine.

**Figure 1.**
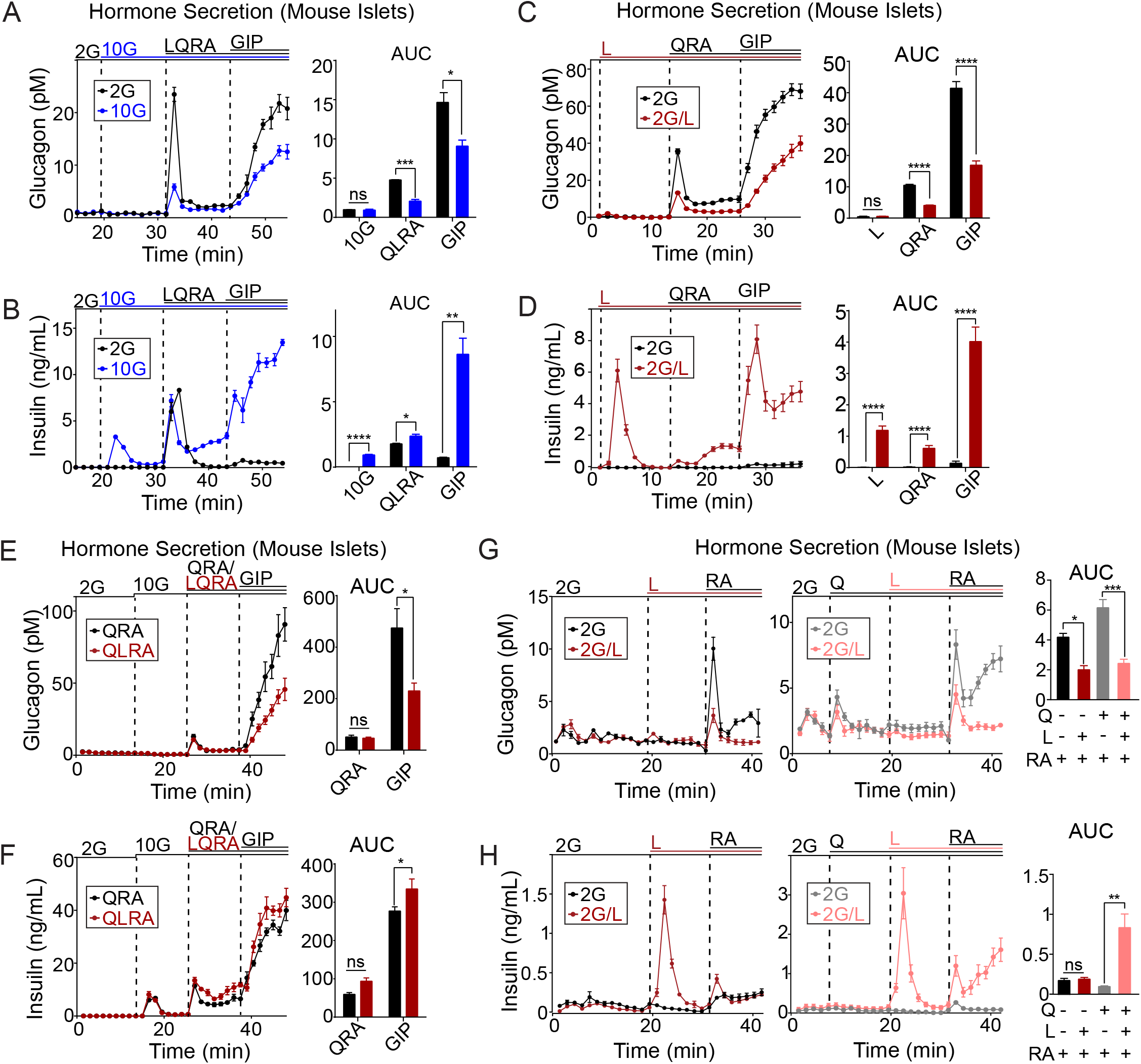
Leucine suppresses glucagon secretion. (A-H) Perifusion assays were used to simultaneously measure glucagon and insulin release from mouse islets stimulated with 2 or 10 mM glucose (2G and 10G), amino acids provided at three times physiological concentration (L, 1.5 mM leucine; Q, 1.8 mM glutamine; R, 0.6 mM arginine; A, 6.3 mM alanine), and GIP (10 nM) as indicated. 90-100 islets/chamber from n = 6 mice/condition (A-D) or n = 3 mice/condition (E-H). Data are shown as mean ± SEM for time traces and area under the curve (AUC) for each pair analyzed by Student’s t-test with *p < 0.05, **p < 0.01, ***p < 0.001, ****p < 0.0001.

Similarly to glucose, leucine strongly suppressed glucagon secretion stimulated by the glucagonotropic amino acid mixture QRA, including when GIP was present (Fig. 1C). Leucine was also sufficient to stimulate insulin secretion, which was further amplified by QRA and GIP (Fig. 1D). Additionally, leucine suppressed GIP-stimulated glucagon secretion and stimulated insulin secretion at 10 mM glucose (Fig. 1E, F). We further tested leucine in the presence of glutamine, which anaplerotically fuels the TCA cycle by maintaining the glutamate pool. While glutamine alone had no effect on insulin or glucagon secretion, it increased glucagon secretion stimulated by RA, and boosted leucine-stimulated insulin secretion (Fig. 1G, H), indicating that glutamine is important for the full secretory response of both α and β-cells. However, leucine was effective at suppressing glucagon secretion whether or not glutamine was present (Fig. 1G, H). Taken together, these findings suggest an important role for leucine in the negative regulation of α-cells at both low and high glucose concentrations.

### Leucine dose-dependently inhibits α-cell cAMP, while having no effect on α-cell Ca^2+^

To determine whether leucine suppresses glucagon secretion through α-cell Ca^2+^ or cAMP, islets were isolated from GcgCre^ERT^:GCaMP6s and GcgCre^ERT^:CAMPER mice, respectively. At low glucose, LQRA stimulated a strong increase in α-cell Ca^2+^, an effect that was due to QRA (Fig. 2A). These findings are consistent with prior reports that arginine and alanine depolarize the α-cell plasma membrane [7,13,43]. Leucine, in contrast, had a negligible effect on α-cell calcium (Fig. 2A), arguing against a Ca^2+^-dependent mechanism for its glucagonostatic effect.

**Figure 2.**
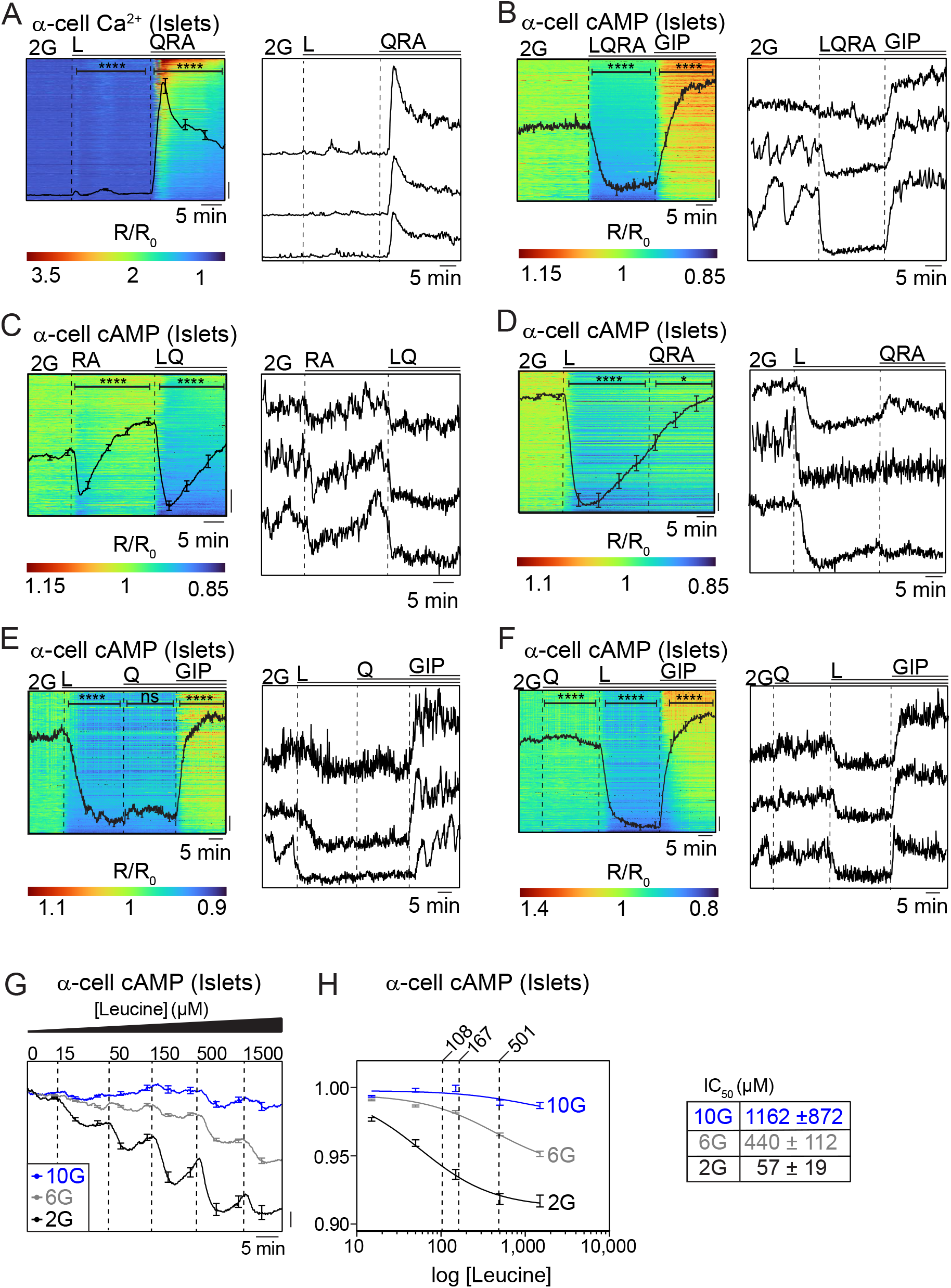
Leucine dose-dependently inhibits α-cell cAMP, while having no effect on α-cell Ca^2+^. (A-F) Measurements of α-cell Ca^2+^ (A) or α-cell cAMP (B-F) from intact mouse islets stimulated with glucose, amino acids, and GIP as in Fig. 1. The left panel shows intact islet averages with individual α-cell responses behind. Representative single-cell traces are shown in the right panel from three α-cells within a representative islet. Data reflect 273-924 single α-cells and 62-131 islets from n = 3 mice/condition. (G and H) Dose-response of α-cell cAMP to leucine (0.015-1.5 mM) quantified from islet averages at 2 mM glucose (2G, *black*), 6 mM glucose (6G, *gray*), or 10 mM glucose (10G, *blue*). Data reflect 68-73 islets from n = 3 mice/condition. The IC_50_ at each level of glucose was calculated using best-fit curve. Data are shown as mean ± SEM for time traces and area under the curve (AUC) for each pair analyzed by Student’s t-test with *p < 0.05, **p < 0.01, ***p < 0.001, ****p < 0.0001.

In islets isolated from GcgCre^ERT^:CAMPER mice, LQRA strongly suppressed α-cell cAMP (Fig. 2B), including when LQRA was delivered at its physiological concentration in the portal circulation [7] (Supplemental Fig. 1A). Single cell analysis (displayed as one cell per row in Fig. 2B) revealed a heterogenous response to LQRA, even within α-cells of the same islet, however cAMP pulsatility was strongly suppressed across the entire population. GIP (10 nM), applied as a positive control that increases α-cell cAMP via the α-cell GIP receptor [7], reversed the effect of LQRA (Fig. 2B). When separated, RA caused a transient decrease in α-cell cAMP, while LQ resulted in a more pronounced cAMP reduction (Fig. 2C). Leucine alone was sufficient to decrease α-cell cAMP and terminate cAMP pulsatility, and the further addition of QRA had no effect (Fig. 2D). The addition of glutamine prior to leucine did not further potentiate the drop in cAMP or change the kinetics of the drop (Fig. 2E and 2F), consistent with the islet perifusion assays (Fig. 1).

Given that leucine reduced α-cell cAMP in the above experiments, we next performed dose-response curves to determine the IC_50_ at varying levels of glucose. Leucine was applied to intact islets between 15 μM and 1.5 mM, which encompasses the fasting and fed concentrations of leucine as measured from the tail vein of mice (108 and 167 μM, respectively) [44,48], as well as the higher postprandial concentration found in the portal blood (501 μM) [7]. As glucose was elevated from 2 to 6 mM, the IC_50_ of leucine increased from 57 ± 19 μM (n = 68 islets) to 440 ± 112 μM (n = 73 islets). In the presence of 10 mM glucose, leucine suppressed α-cell cAMP starting at 150 μM, and the IC_50_ increased to 1162 ± 872 μM (n = 71 islets) (Fig. 2G and 2H). Combined, these data emphasize the physiological action of leucine to suppress α-cell cAMP across the physiological range of glucose.

### Leucine and glucose regulate α-cell cAMP and glucagon secretion independently of Sstr2

In mouse islets, prior studies with the Sstr2 antagonist CYN154806 (CYN) demonstrate an important role for tonic somatostatin regulation of the α-cell [28,48,49]. In the presence of CYN, high glucose stimulated insulin secretion and reduced LQRA-stimulated glucagon secretion (Fig. 3A, B). Notably, high glucose failed to dampen GIP-stimulated glucagon secretion in the presence of CYN (Fig. 3A), suggesting an interaction between Sstr2 and GIP receptor signaling. Like glucose, leucine effectively suppressed RA-stimulated glucagon secretion in the presence of CYN (Fig. 3C). Similar results were obtained in the presence of glutamine, which increased both glucagon and insulin release (Fig. 3C, D).

**Figure 3.**
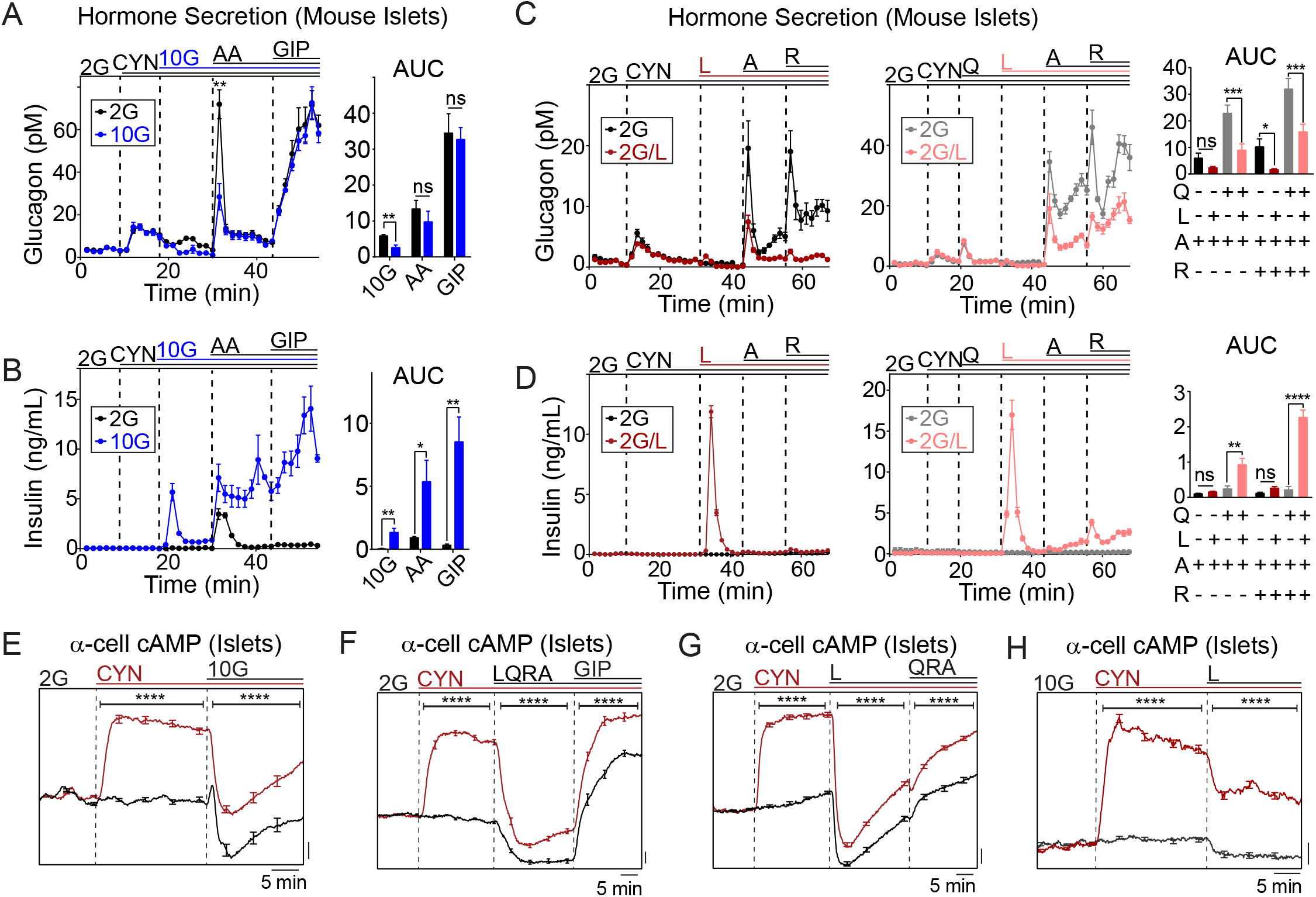
Leucine and glucose regulate α-cell cAMP and glucagon secretion independently of Sstr2. (A-D) Perifusion assays of mouse islets stimulated with glucose, amino acids, and GIP as in Fig. 1, with the inclusion of 500 nM SSTR2 antagonist CYN154806 (CYN) as indicated. 90-100 islets/chamber from n = 3 mice per group. (E-H) Epifluorescence microscopy of α-cell cAMP in intact mouse islets stimulated with glucose, amino acids, and GIP as in Fig. 1. Data reflect 43-91 islets from n = 3 mice/group. Data are shown as mean ± SEM for time traces and AUC for each condition with *p < 0.05, **p < 0.01, ***p < 0.001, ****p < 0.0001 by Student’s t test (A,B,E-H) or one-way ANOVA (C and D).

In parallel with the glucagon secretion measurements, Sstr2 inhibition strongly increased α-cell cAMP in the presence of 2 mM glucose (Fig. 3E). The further addition of glucose effectively lowered α-cell cAMP in the presence of CYN, consistent with δ-cell independent regulation of α-cell cAMP. Similarly, LQRA and leucine decreased α-cell cAMP with CYN present at both low and high glucose (Fig. 3F-H). Collectively, these data are consistent with both Sstr2-dependent and Sstr2-independent mechanisms for leucine suppression of α-cell cAMP and glucagon secretion.

### Leucine modulates α -cell cAMP via α -cell intrinsic and islet paracrine signaling

To test for an α-cell intrinsic effect of leucine in intact islets, we used the K_ATP_ channel opener diazoxide (200 μM) to block Ca^2+^-dependent exocytosis from α, β, and δ-cells [14,50,51]. Diazoxide increased α-cell cAMP to a similar extent as CYN (Fig. 4A). LQRA effectively lowered α-cell cAMP in the presence of diazoxide, but not to the same level as in its absence, suggesting an α-cell intrinsic mechanism in addition to a paracrine effect. GIP was effective at raising α-cell cAMP in the presence of diazoxide (Fig. 4A), consistent with a direct effect on the α-cell GIP receptor [7]. RA induced only a transient decrease in cAMP, although this effect was reversed by diazoxide (Fig. 4B) and may be Ca^2+^-dependent. Similar effects were observed at physiological levels of RA (Supplemental Fig. 1B). Leucine, alone or in combination with glutamine, strongly suppressed α-cell cAMP in the presence of diazoxide (Fig. 4B-E). Leucine had no impact on α-cell Ca^2+^ in the presence of diazoxide, which blocked LQRA-stimulated Ca^2+^ influx (Fig. 4F). Finally, we used islet dispersion to eliminate paracrine and juxtacrine signaling. In isolated α-cells identified by the expression of CAMPER, leucine inhibited α-cell cAMP, with no further effect of QRA (Fig. 4G). Taken together with the CYN and diazoxide experiments in intact islets, these findings in dispersed α-cells confirm a direct, suppressive effect of leucine on α-cell cAMP.

**Figure 4.**
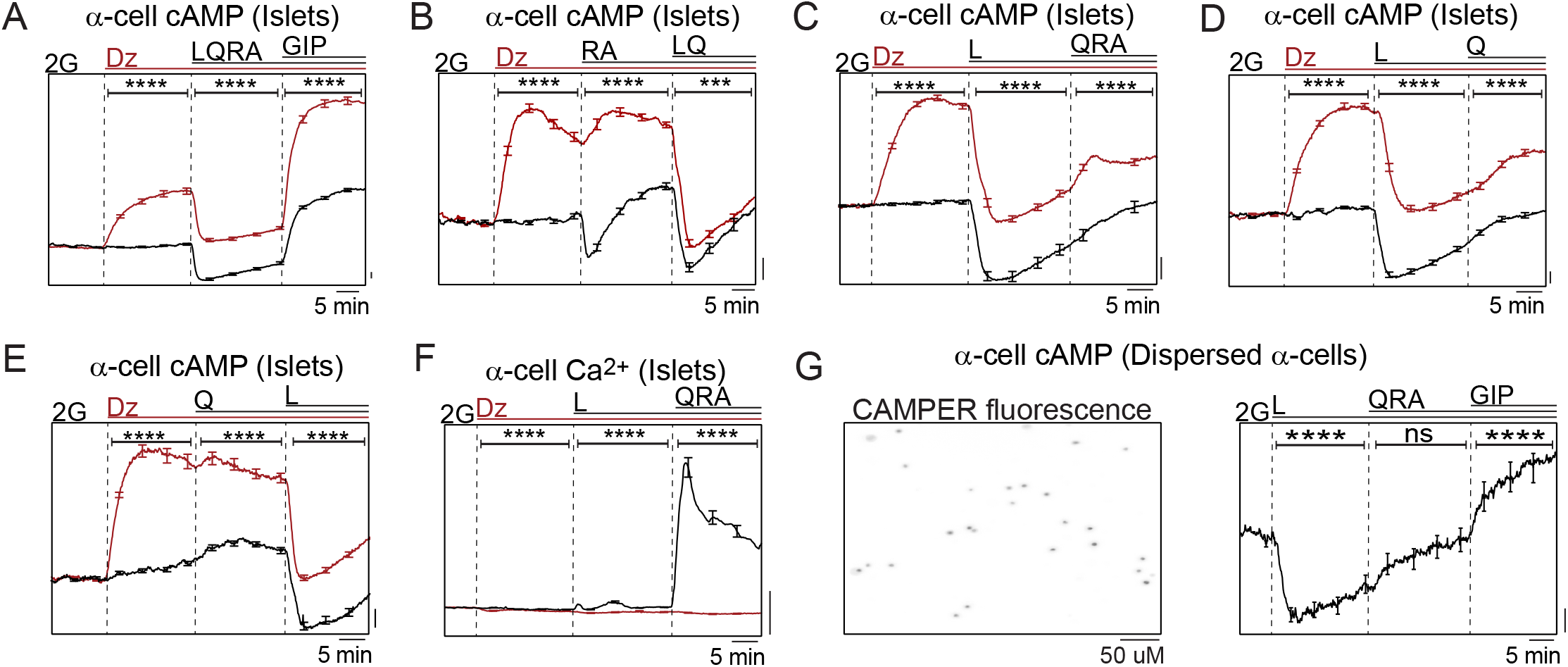
Leucine modulates α-cell cAMP via both α-cell-intrinsic and islet paracrine signaling. Epifluorescence microscopy α-cell cAMP in intact islets (A-E), α-cell Ca^2+^ in intact islets (F) stimulated with glucose, amino acids, and GIP as in Fig. 1, with the inclusion of 200 μM diazoxide (Dz) as indicated. Data reflect 50-94 islets from n = 3 mice/group. (G) Representative image of dispersed α-cells expressing CAMPER (*left*), followed by cAMP time courses (*right*). Data reflect 141 α-cells from n = 3 mice. Data are shown as mean ± SEM for each condition with ***p < 0.001, ****p < 0.0001 by Student’s t test.

### Leucine suppresses α-cell cAMP by stimulating mitochondrial anaplerosis

The data above indicates that leucine mimics glucose regulation of α-cell cAMP and glucagon secretion. In the presence of diazoxide, 10 mM glucose suppressed α-cell cAMP to the same extent as 1.5 mM leucine, which had no further effect when added after glucose (Figs. 5A and B). These findings are consistent with a common, mitochondrial-dependent mode of action. If the mechanism of cAMP suppression involved acetyl-CoA generation, and activation of the TCA cycle, 10 mM glucose would be expected to have a much stronger effect on cAMP than 1.5 mM leucine. Instead, the higher potency of leucine suggests a mechanism dependent on anaplerosis. Glucose carbons anaplerotically enter the TCA cycle through pyruvate carboxylase, which has low activity in α-cells [52], while leucine allosterically activates glutamate dehydrogenase to increase glutamate conversion to αLJketoglutarate [53]. The application of α-ketoisocaproate (KIC), which can be transaminated to leucine [54,55], had a similar effect on α-cell cAMP to glucose, a further indication that mitochondrial fuels are sufficient to regulate α-cell cAMP (Fig. 5C). Methyl-succinate, a complex II fuel that only weakly supports anaplerosis, was far less effective at lowering α-cell cAMP, which was subsequently reduced by leucine addition (Fig. 5D). To directly test whether leucine regulates α-cell cAMP through a mechanism dependent on mitochondrial anaplerosis, we applied 2-aminobicyclo(2,2,1)heptane-2-carboxylic acid (BCH), a non-metabolizable leucine analog that stimulates anaplerosis by activating glutamate dehydrogenase. BCH strongly reduced α-cell cAMP, indicating that glutamate dehydrogenase activation rather than acetyl-CoA generation underlies leucine suppression of cAMP (Fig. 5E). Collectively, these data suggest that mitochondrial anaplerosis facilitates α-cell cAMP regulation by glucose and leucine.

**Figure 5.**
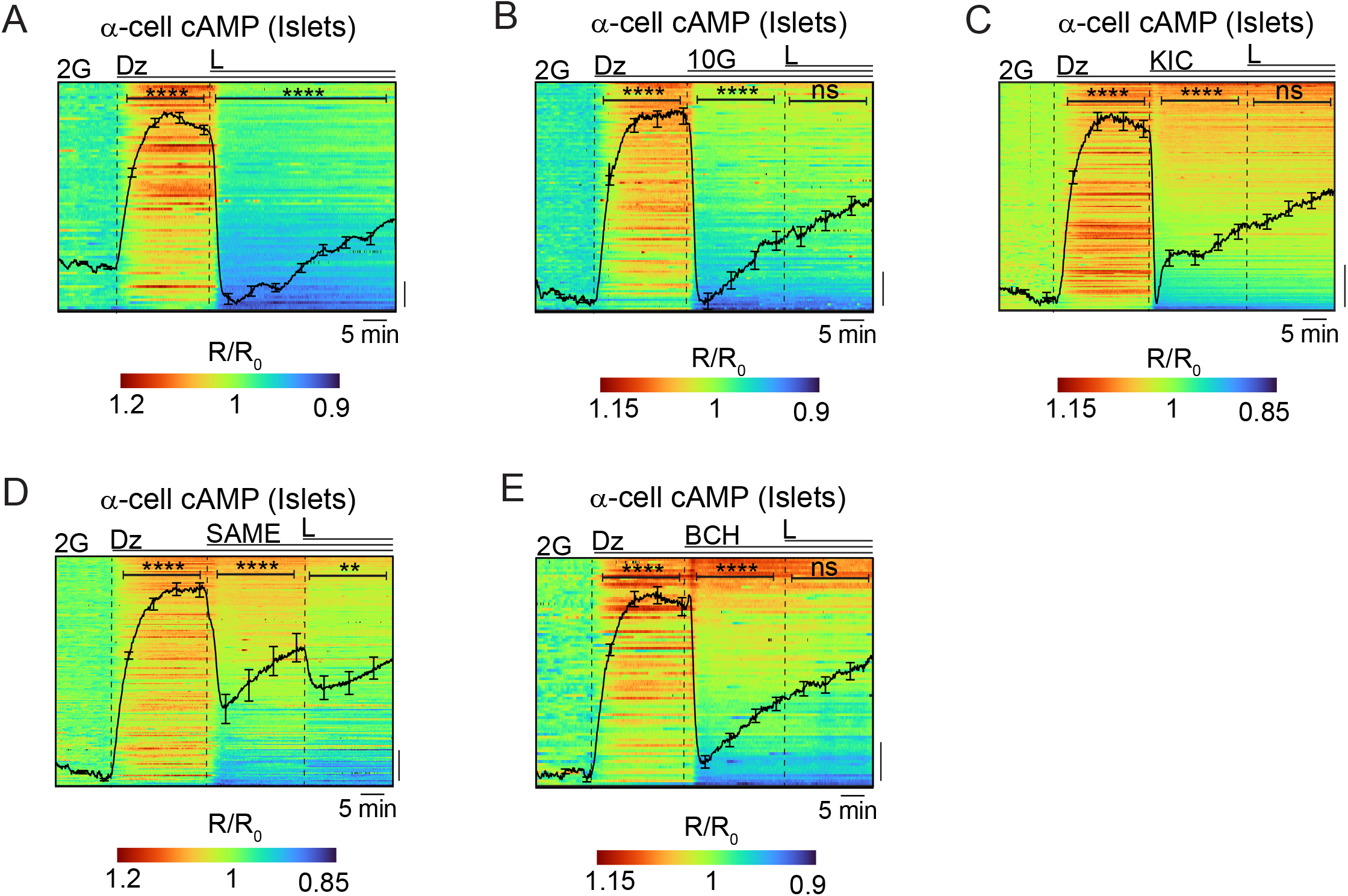
Leucine suppresses α-cell cAMP by stimulating mitochondrial anaplerosis. (A-E) Epifluorescence microscopy of α-cell cAMP in intact mouse islets represented as islet averages, with kymographs of individual α-cell responses behind. Islets were imaged in 2 mM glucose (2G) and 200 μM diazoxide (Dz), with, 1.5 mM leucine (L), 10 mM glucose (10G), 10 mM α-ketoisocaproate (KIC), 10 mM methyl-succinate (SAME), or 10 mM 2-aminobicyclo(2,2,1)heptane-2-carboxylic acid (BCH) added as indicated. Data reflect 82-177 single α-cells and 39-65 islets from n = 3 mice/condition. Data are shown as mean ± SEM for each condition with **p < 0.01, ****p < 0.0001 by Student’s t test.

### Leucine regulates amino acid-stimulated glucagon release from primary human islets

Primary human islets were used in perifusion assays to test the effect of leucine on glucagon and insulin secretion. Matching its effects on mouse islets, leucine suppressed QRA-stimulated glucagon secretion in all three donors tested (Fig. 6). On a technical note, α-cells from human islets are more sensitive than mouse islets to flow changes, which in three out of four preparations resulted in artifactual increases in glucagon secretion that occur during fraction collector plate changes. This spurious effect is most apparent when the control trace, which did not receive leucine, exhibited an increase in glucagon but not insulin. Nonetheless, the suppressive effect of leucine on glucagon secretion is consistent with the effect in mouse islets.

**Figure 6.**
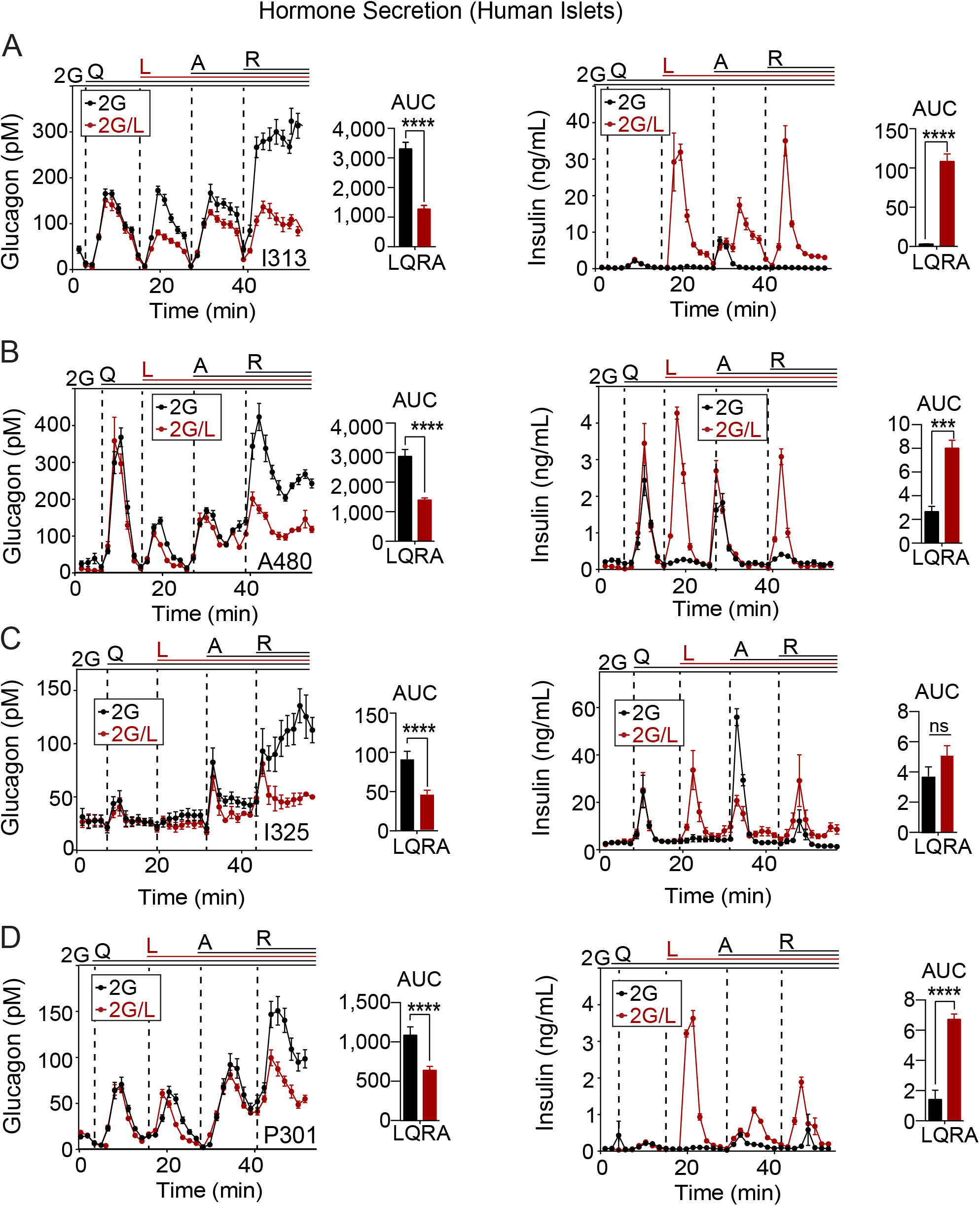
Leucine regulates glucagon secretion in primary human islets. (A-D) Perifusion assays were used to simultaneously measure insulin and glucagon secretion from human islets stimulated with glucose and amino acids as in Fig. 1. 90-100/islets chamber x 12 chambers per donor from n = 4 human donors. Data are shown as mean ± SEM for time traces and AUC for each condition with *p < 0.05, **p < 0.01, ***p < 0.001, ****p < 0.0001 by Student’s t test.

## Discussion

Our findings reveal a role for leucine in the regulation of α-cell cAMP and glucagon secretion, particularly under hypoglycemic and euglycemic conditions. Like glucose, leucine exerts a direct suppressive effect on α-cell cAMP, in addition to stimulating inhibitory paracrine signals from β/δ-cells. Although the IC_50_ of leucine increases with glucose, it remains well-matched to the fasting and fed levels of leucine in the portal circulation [7]. While these *ex vivo* studies may not precisely match the *in vivo* situation, they suggest that leucine levels in the blood help to set the tone of glucagon secretion in the fasted state, while glucose and leucine share control over glucagon secretion in the fed state. Consistently, leucine was effective at limiting GIP-stimulated glucagon secretion, which is important for boosting β-cell function when glucose is elevated [5,7,8,56–58]. Leucine suppressed α-cell cAMP independently of Ca^2+^, K_ATP_ channel activity, and paracrine/juxtacrine signaling, as shown by the use of pharmacological paracrine blockers within intact islets, and by dispersed α-cells. However, the nadir in α-cell cAMP was blunted by pharmacological paracrine blockers, consistent with a dual mechanism of intrinsic and extrinsic α-cell regulation by leucine. Finally, we provide evidence that α-cell intrinsic cAMP regulation is dependent upon mitochondrial anaplerosis.

When applied by themselves, neither glucose nor leucine strongly impacted glucagon secretion, despite reducing α-cell cAMP. However, both fuels are easily seen as negative regulators of α-cell tone when electrogenic amino acids such as alanine and arginine are present to stimulate Ca^2+^ influx. These findings support a general model for islet cell regulation in which depolarization controls the timing of hormone release, while cAMP sets the magnitude of secretion [6,59]. For example, the incretin hormones GLP-1 and GIP can raise β-cell cAMP, but their insulinotropic effect requires a depolarizing stimulus (from glucose); a similar model has been proposed for UCN3 potentiation of somatostatin release from δ-cells [60]. Layered on top of this basic framework is the concept that a heterogeneous population of α-cells, each with its own distinct nutrient response, collectively sets glucagon output from the islet, as revealed by our single cell analyses of both Ca^2+^ and cAMP. Although strong perturbations in the α-cell population have been observed in type 2 diabetes with glucose as a fuel [39], leucine remains untested.

Metabolic fuels including leucine, BCH, KIC, and glucose were each sufficient to suppress α-cell cAMP with the Ca^2+^-mediated exocytosis blocker diazoxide present, implying an intrinsic effect that is dependent on mitochondrial metabolism. Since diazoxide lowers α-cell Ca^2+^, these experiments also argue against a dominant role of Ca^2+^-dependent phosphodiesterases or adenylate cyclases as a central mechanism by which these fuels control α-cell cAMP. Although most of these mitochondrial fuels stimulate both oxidative and anaplerotic pathways, multiple lines of evidence are consistent with a mechanism dependent on mitochondrial anaplerosis. The complex II fuel methyl-succinate, once converted to succinate, must exchange for malate upon entering the mitochondria (given the absence of malic enzyme in mouse islets that is needed to make pyruvate [61]). Consequently succinate can only indirectly stimulate anaplerosis by raising the mitochondrial voltage, which is expected to slow the TCA cycle and raise acetyl-CoA to activate anaplerotic pyruvate consumption from glucose by pyruvate carboxylase [62]. Accordingly, methyl-succinate reduced cAMP to a much lesser extent than leucine. In comparison, the glutamate dehydrogenase activator BCH, a purely anaplerotic fuel that is not metabolized to acetyl-CoA, suppressed α-cell cAMP and was not further impacted by leucine addition. The fact that BCH and glucose have the same suppressive effect on cAMP, but stimulate anaplerosis via different mechanisms (i.e., glutamate dehydrogenase and pyruvate carboxylase), argues for a general role of anaplerosis in controlling cAMP production and setting α-cell tone.

The much higher potency of leucine (μM) versus glucose (mM) for suppressing α-cell cAMP may be explained by the low level of α-cell pyruvate carboxylase activity [52], which mediates anaplerosis of glucose carbons. An alternative, and highly complementary explanation, is that the high level of lactate dehydrogenase in α-cells [38,52,63] shunts glucose-dependent pyruvate away from α-cell mitochondria, such that glycolysis is poorly coupled to α-cell oxidative phosphorylation [25,52,64] (i.e., α-cells favor aerobic glycolysis, just the opposite of β-cells [65]). Both of these α-cell specializations would also limit the ability of leucine to stimulate pyruvate anaplerosis via the phosphoenolpyruvate cycle, which is dependent on efficient mitochondrial pyruvate uptake as well as pyruvate metabolism via pyruvate carboxylase [62,66,67]. Consequently, it seems likely that leucine impacts α-cell cAMP primarily via glutamate dehydrogenase, while glucose is forced to work less efficiently via pyruvate carboxylase.

Integrating our work with previous studies, a model of α-cell metabolic regulation is beginning to take shape. Under hypoglycemic conditions, when glucagon secretion is maximal, α-cell mitochondria rely on fatty acid oxidation for ATP production [40,41]; under these conditions, cAMP and Ca^2+^ are elevated [3,28,41]. Glucose metabolism, by stimulating citrate cataplerosis, generates malonyl-CoA, which blocks mitochondrial fatty acid import via CPT1 inhibition [40,41]. Although blockade of β-oxidation might be expected to reinforce glucose utilization in the TCA cycle and boost cytosolic ATP/ADP, as it does in β-cells [52], α-cell mitochondria oxidize very little glucose and exhibit low levels of glucose-stimulated ATP production [25,52,64]. A recent study further indicates that glucose-dependent inhibition of fatty acid oxidation *reduces* cytosolic ATP/ADP in α-cells [41]. Glucose-dependent suppression of mitochondrial ATP production, while paradoxical, provides a compelling explanation for the ability of both glucose and leucine to suppress cytosolic cAMP and glucagon secretion, as demonstrated here. Unfortunately, we do not yet know whether leucine mimics the ability of glucose to shut down fatty oxidation and reduce cytosolic ATP/ADP. However, the available evidence suggests that α-cell mitochondria favor anaplerosis-cataplerosis over oxidation [25,41,52,64], (i.e., Mito_Cat_ is favored over Mito_Ox_ [68]), thus allowing both glucose and leucine to limit α-cell tone by reducing metabolism-dependent cAMP production. This model awaits further testing.

### Limitations and technical considerations

An potential limitation is that our studies of nutrient-regulated cytosolic cAMP generation may not account for localized, plasma-membrane compartmentalized ATP and cAMP signaling in α-cells [17,28,38]. Additionally, anaplerosis-cataplerosis is likely not the only possible mechanism for cAMP regulation, as mTORC signaling may also play a role in amino acid sensing [69], at least chronically [70]. We attempted glutamine deprivation as a method to prevent leucine-stimulated anaplerosis, a technique used previously in β-cells to identify the function of glutamate dehydrogenase [53], including mutations that cause hyperinsulinemic-hyperammonemia [71]. However, leucine effectively suppressed cAMP and glucagon secretion in the absence of glutamine (plausibly because glutamate is not limiting). Nonetheless, these experiments reveal that glutamine is essential for the full secretory response of both α and β-cells in response to glucose and/or amino acid stimulation. In future studies, we recommend the inclusion of 0.5 mM glutamine (i.e., the physiological level in the portal blood [7]) in all solutions in which human or mouse islets are examined *ex vivo*.

As described in the last section of results, we observed a spurious increase in glucagon secretion during fraction collector plate changes, particularly human islets studied at low glucose. Since our leucine experiments were controlled, these artifacts did not affect our conclusions. However, it is important that solution flow remains uninterrupted for perifusion measurements of glucagon secretion. This can be achieved using the BioRep Peri-5 with a software patch that is now available following discussions with the manufacturer. Finally, as detailed in the methods, we successfully performed studies with GcgCre^ERT^ mice [72] without using tamoxifen. While it may be possible to perform tamoxifen-inducible studies if GcgCre^ERT^ is present in the sire, spurious GCaMP6s fluorescence was observed in the absence of tamoxifen (or 4-hydroxytamoxifen) when GcgCre^ERT^ was present in the dam (Suppl. Table 2). This is potentially an advantage since tamoxifen itself is not inert [73,74]. We achieved 89% knockdown of *Pck2* mRNA in FACS-sorted α-cells from 12-week-old GcgCre^ERT^:GCaMP6s:Pck2^f/f^ mice by comparison to GcgCre^ERT^:GCaMP6s:Pck2^wt/wt^ littermate controls; here, 4-hydroxytamoxifen was included in the islet culture media (unpublished observations). Investigators utilizing the tamoxifen-inducible GcgCre^ERT^ system should tailor the breeding strategy to the experimental goals.

## Acknowledgements

This work was supported in part by the United States Department of Veterans Affairs Biomedical Laboratory Research and Development Service (I01BX005113 to MJM). The Merrins laboratory also gratefully acknowledges support from the NIH/NIDDK (R01DK113103 and R01DK127637). ERK received a predoctoral fellowship from the NIH/NIDDK (F31DK134171). Human pancreatic islets were provided, in part, by the NIDDK-funded Integrated Islet Distribution Program (IIDP) (RRID:SCR_014387) at City of Hope (UC4DK098085) and the JDRF-funded IIDP Islet Award Initiative. This work utilized facilities and resources from the William S Middleton Memorial Veterans Hospital and does not represent the views of the Department of Veterans Affairs or the United States Government.

## Methods

### Mice and Islet Isolations

GcgCre^ERT^ mice [72] (Jax 030346) were crossed with GCaMP6s mice (Jax 028866), a Cre-dependent Ca^2+^ indicator strain [75], and CAMPER mice (Jax 032205), a Cre-dependent cAMP indicator strain [76]. As in our previous study [7], the mice were not treated with tamoxifen (additional details are provided in the next section). Mice were sacrificed by CO_2_ asphyxiation followed by cervical dislocation at 12-15 weeks of age, and islets were isolated as detailed in [77]. Islets were cultured in RPMI-1640 supplemented with 10% (v/v) fetal bovine serum (ThermoFisher A31605), 10,000 units/mL penicillin and 10,000 mg/mL streptomycin (Fisher Scientific). All procedures involving animals were approved by the Institutional Animal Care and Use Committees of the William S. Middleton Memorial Veterans Hospital, and followed the NIH Guide for the Care and Use of Laboratory Animals.

### *Imaging of* α*-cell cAMP and Ca*^*2+*^

Islets isolated from GcgCre^ERT^:GCaMP6s mice were cultured overnight in the presence of 100 nM 4-hydroxytamoxifen (Sigma) and imaged three days post isolation. Notably, we found that 4-hydroxytamoxifen was not necessary for GCaMP6s expression provided that GcgCre^ERT^ was present in the dam (Suppl. Table 2). Leveraging this breeding strategy to include GcgCre^ERT^ in both the sire and dam, islets isolated from GcgCre^ERT^:CAMPER offspring expressed the reporter at maximal levels one day post isoltation in the absence of 4-hydroxytamoxifen, which was not used. In both cases, the leaky reporter expression was due to spurious Cre-dependent recombination, since expression of GCaMP6s and CAMPER was not observed in the absence GcgCre^ERT^. Isolated islets were imaged in an RC-41LP imaging chamber inserted in a QE-1 chamber holder (Warner Instruments) mounted on a Nikon Ti microscope equipped with a 10×/0.5NA SuperFluor objective (Nikon Instruments). The chamber was perfused with a standard bath solution containing 137 mM NaCl, 5.6 mM KCl, 1.2 mM MgCl_2_, 0.5 mM NaH_2_PO_4_, 4.2 mM NaHCO_3_, 10 mM HEPES, and 2.6 mM CaCl_2_ with varying concentrations of glucose (2, 6, or 10 mM) at pH 7.4. Glutamine (0.6-1.8 mM), leucine (0.015-1.5 mM), arginine (0.2-0.6 mM), alanine (2.1-6.3 mM), GIP (10 nM), BCH (10 mM), KIC (10 mM), methyl-succinate (10 mM), diazoxide (200 μM), and CYN154806 (500 nM) were added as indicated in the figure legends. The flow rate was maintained at 0.25-4 mL/min using a feedback-controlled flow cell (Fluigent MCFS-EZ) and temperature was maintained at 33°C using solution and chamber heaters (Warner Instruments). LED excitation for CAMPER was provided by either Semrock 434/21, 5% output from an AURA light engine (Lumencor) or 434/21 5% output from a Spectra III light engine (Lumencor) with a Semrock dichroic FF459/526/596-Di01 and emission filters Semrock FF02-470/30 and FF01-542/27. Fluorescence emission for CAMPER was collected every 6 seconds with a Prime 95B back-illuminated sCMOS camera (Photometrics) with 2x2 binning. LED excitation for GCaMP6s was provided by a SOLA SE II 365 (Lumencor) set to 20% output and an inline 25% neutral density filter (Nikon ND4). Fluorescence emission was collected with a Hamamatsu ORCA-Flash4.0 V2 Digital CMOS camera at every 6 seconds. Excitation (x) and emission (m) filter were used in combination with a FF458-Di02 dichroic (Semrock): GCaMP6s, ET500/20x, ET535/30m. A single region of interest was used to quantify the average response of each islet or dispersed cell using Nikon Elements, with cAMP levels reported as the CAMPER emission ratio R470/542 normalized to the initial condition (R/R_o_), and Ca^2+^ levels reported as the GCaMP6s intensity normalized to the initial condition (F/F_o_). Single α-cell quantification of CAMPER and GCaMP6s was performed with the Spots tool using Imaris software (Bitplane).

### Human Islets

Human islets from normal donors were obtained from the Alberta Diabetes Institute Islet Core, Prodo Labs, and Integrated Islet Distribution Program. The age, sex, body mass index, and %HbA1c of each donor are provided in Supplementary Table S1. Islets were cultured in Prodo Islet Media [Pim(R)] complete media (Prodo Laboratories) for a period of 24-48 hours after being received. This study used data from the Organ Procurement and Transplantation Network (OPTN). The OPTN data system includes data on all donor, wait-listed candidates, and transplant recipients in the US, submitted by the members of the Organ Procurement and Transplantation Network (OPTN). The Health Resources and Services Administration (HRSA), U.S. Department of Health and Human Services provides oversight to the activities of the OPTN contractor. The data reported here have been supplied by UNOS as the contractor for the Organ Procurement and Transplantation Network (OPTN). The interpretation and reporting of these data are the responsibility of the author(s) and in no way should be seen as an official policy of or interpretation by the OPTN or the U.S. Government.

### Islet perifusion assays

For each assay, islets from 12 mice or from a single human donor were pooled and then placed into a 12-channel perifusion system (BioRep Peri-5) with 90-100 islets/chamber. Islets are equilibrated at 2 mM glucose for 48 minutes at 37ºC in KRPH buffer (in mM: 140 NaCl, 4.7 KCl, 1.5 CaCl_2_, 1 NaH_2_PO_4_, 1 MgSO_4_, 2 NaHCO_3_, 5 HEPES, 0.1% BSA, pH 7.4) with 100 μL Bio-Gel P-4 Media (1504124) before stimulation as described in the figure legends. Insulin and glucagon secretion were assayed using Lumit Insulin Immunoassay (Promega CS3037A01) and Lumit Glucagon Immunoassay (Promega W8022) and measured using a TECAN Spark plate reader.

### Statistics

All data are presented as means ± SEM. For perifusion assays, AUCs were analyzed using an unpaired t-test for 2 conditions while 1-way ANOVA was used when comparing 4 conditions. Imaging data were normalized to the control condition. Unpaired t-tests were used when comparing the AUCs between control and experimental conditions. Paired t-tests were used when comparing each imaging condition to the preceding (control) condition (note that ANOVA is inappropriate in this case because only 2 groups can be compared in a controlled fashion).

## Figure Legends

**Supplemental Figure 1.** (A) Epifluorescence microscopy of α-cell cAMP in intact mouse islets stimulated with 2 mM glucose (2G), amino acids provided at physiological concentration (L, 0.5 mM leucine; Q, 0.6 mM glutamine; R, 0.2 mM arginine; A, 2.1 mM alanine), and GIP (10 nM) as indicated. Data reflect 147 islets from n = 4 mice. (B) Epifluorescence microscopy of α-cell cAMP in intact mouse islets stimulated with 2 mM glucose (2G), amino acids provided at physiological concentration (L, 0.5 mM leucine; Q, 0.6 mM glutamine; R, 0.2 mM arginine; A, 2.1 mM alanine), and 200 μM diazoxide (Dz) as indicated. Data reflect 147 islets from n = 4 mice. Data are shown as mean ± SEM for each condition with ****p < 0.0001 by Student’s t test.

**Supplemental Table 1.**
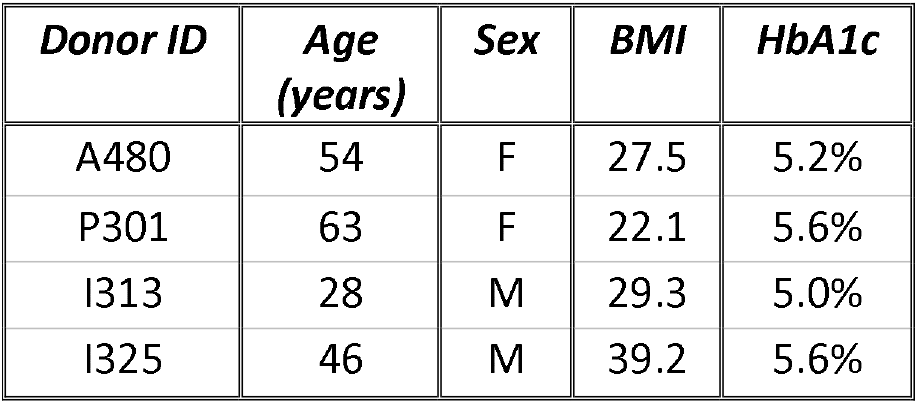
Summary characteristics of human islet donors. BMI, body mass index; HbA1c, glycated hemoglobin.

**Supplemental Table 2.**
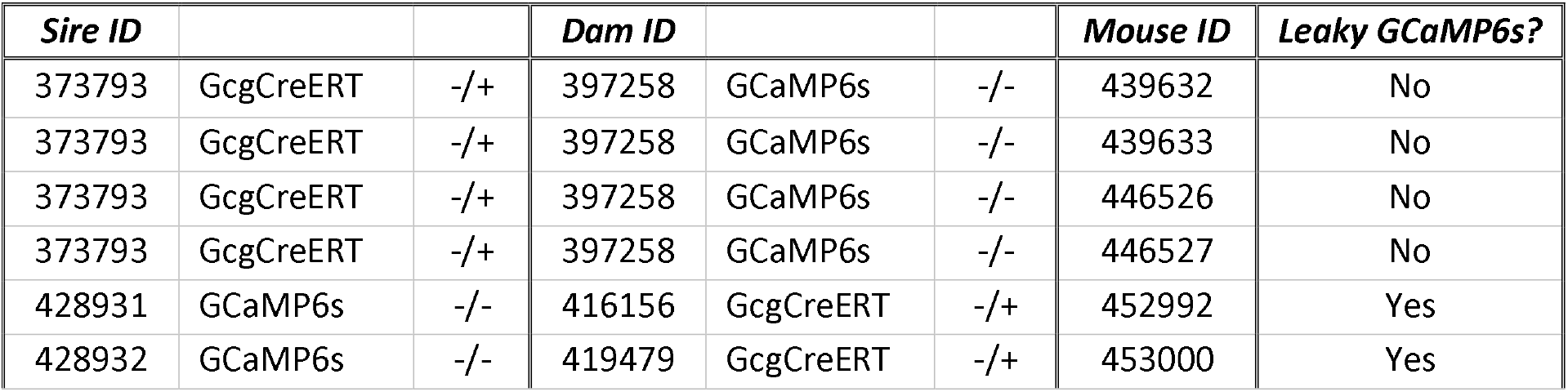
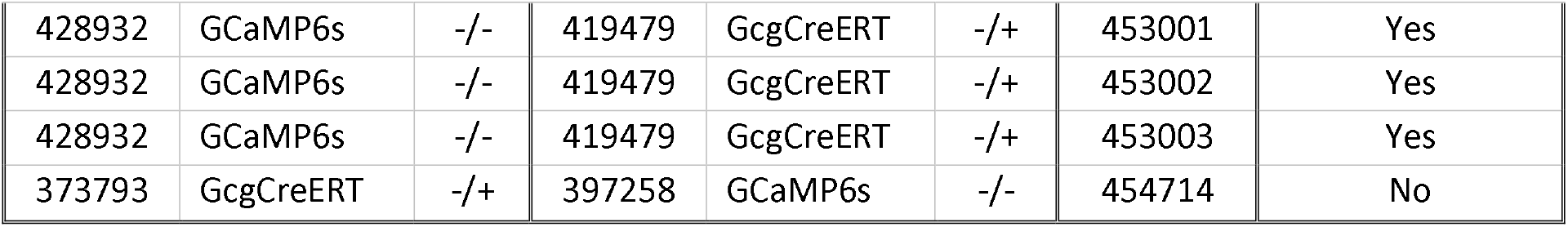
Qualitative determination of reporter expression observed in the islets from GcgCre^ERT^:GCaMP6s mice. Spurious expression of GCaMP6s was observed, *i*.*e*. in the absence of tamoxifen or 4-hydroxytamoxifen, when GcgCre^ERT^ was present in the dam (n = 10 mice from 4 breeding pairs). No expression of GCaMP6s was observed in the absence of GcgCre^ERT^.

## References

[1] Wewer Albrechtsen, N.J., Holst, J.J., Cherrington, A.D., Finan, B., Gluud, L.L., Dean, E.D., et al., 2023. 100 years of glucagon and 100 more. Diabetologia 66(8): 1378–94, Doi: 10.1007/s00125-023-05947-y.

[2] Dean, E.D., 2020. A Primary Role for α-Cells as Amino Acid Sensors. Diabetes 69(4): 542–9, Doi: 10.2337/dbi19-0021.

[3] MacDonald, P.E., Rorsman, P., 2023. Metabolic Messengers: glucagon. Nature Metabolism 5(2): 186–92, Doi: 10.1038/s42255-022-00725-3.

[4] Murlin, J.R., Clough, H.D., Gibbs, C.B.F., Stokes, A.M., 1923. AQUEOUS EXTRACTS OF PANCREAS. Journal of Biological Chemistry 56(1): 253–96, Doi: 10.1016/S0021-9258(18)85619-8.

[5] Capozzi, M.E., Wait, J.B., Koech, J., Gordon, A.N., Coch, R.W., Svendsen, B., et al., 2019. Glucagon lowers glycemia when β cells are active. JCI Insight 4(16): e129954, Doi: 10.1172/jci.insight.129954.

[6] Capozzi, M.E., Svendsen, B., Encisco, S.E., Lewandowski, S.L., Martin, M.D., Lin, H., et al., 2019. β Cell tone is defined by proglucagon peptides through cAMP signaling. JCI Insight 4(5): e126742, Doi: 10.1172/jci.insight.126742.

[7] El, K., Gray, S.M., Capozzi, M.E., Knuth, E.R., Jin, E., Svendsen, B., et al., 2021. GIP mediates the incretin effect and glucose tolerance by dual actions on α cells and β cells. Science Advances 7(11): eabf1948, Doi: 10.1126/sciadv.abf1948.

[8] Zhu, L., Dattaroy, D., Pham, J., Wang, L., Barella, L.F., Cui, Y., et al., 2019. Intraislet glucagon signaling is critical for maintaining glucose homeostasis. JCI Insight 4(10): e127994, Doi: 10.1172/jci.insight.127994.

[9] Prentki, M., 2006. Islet cell failure in type 2 diabetes. Journal of Clinical Investigation 116(7): 1802–12, Doi: 10.1172/JCI29103.

[10] D’Alessio, D., 2011. The role of dysregulated glucagon secretion in type 2 diabetes. Diabetes, Obesity and Metabolism 13: 126–32, Doi: 10.1111/j.1463-1326.2011.01449.x.

[11] Gilon, P., 2020. The Role of α-Cells in Islet Function and Glucose Homeostasis in Health and Type 2 Diabetes. Journal of Molecular Biology 432(5): 1367–94, Doi: 10.1016/j.jmb.2020.01.004.

[12] Zhang, Q., Ramracheya, R., Lahmann, C., Tarasov, A., Bengtsson, M., Braha, O., et al., 2013. Role of KATP Channels in Glucose-Regulated Glucagon Secretion and Impaired Counterregulation in Type 2 Diabetes. Cell Metabolism 18(6): 871–82, Doi: 10.1016/j.cmet.2013.10.014.

[13] Rorsman, P., Hellman, B., 1988. Voltage-activated currents in guinea pig pancreatic alpha 2 cells. Evidence for Ca2+-dependent action potentials. Journal of General Physiology 91(2): 223–42, Doi: 10.1085/jgp.91.2.223.

[14] MacDonald, P.E., Marinis, Y.Z.D., Ramracheya, R., Salehi, A., Ma, X., Johnson, P.R.V., et al., 2007. A KATP Channel-Dependent Pathway within α Cells Regulates Glucagon Release from Both Rodent and Human Islets of Langerhans. PLoS Biology 5(6): e143, Doi: 10.1371/journal.pbio.0050143.

[15] Gromada, J., Bokvist, K., Ding, W.-G., Barg, S., Buschard, K., Renström, E., et al., 1997. Adrenaline Stimulates Glucagon Secretion in Pancreatic A-Cells by Increasing the Ca2+ Current and the Number of Granules Close to the L-Type Ca2+ Channels. Journal of General Physiology 110(3): 217–28, Doi: 10.1085/jgp.110.3.217.

[16] Barg, S., Galvanovskis, J., Göpel, S.O., Rorsman, P., Eliasson, L., 2000. Tight coupling between electrical activity and exocytosis in mouse glucagon-secreting alpha-cells. Diabetes 49(9): 1500–10, Doi: 10.2337/diabetes.49.9.1500.

[17] Li, J., Yu, Q., Ahooghalandari, P., Gribble, F.M., Reimann, F., Tengholm, A., et al., 2015. Submembrane ATP and Ca ^2+^ kinetics in α-cells: unexpected signaling for glucagon secretion. The FASEB Journal 29(8): 3379–88, Doi: 10.1096/fj.14-265918.

[18] Le Marchand, S.J., Piston, D.W., 2012. Glucose Decouples Intracellular Ca2+ Activity from Glucagon Secretion in Mouse Pancreatic Islet Alpha-Cells. PLoS ONE 7(10): e47084, Doi: 10.1371/journal.pone.0047084.

[19] Le Marchand, S.J., Piston, D.W., 2010. Glucose Suppression of Glucagon Secretion. Journal of Biological Chemistry 285(19): 14389–98, Doi: 10.1074/jbc.M109.069195.

[20] Lavagnino, Z., Dwight, J., Ustione, A., Nguyen, T.-U., Tkaczyk, T.S., Piston, D.W., 2016. Snapshot Hyperspectral Light-Sheet Imaging of Signal Transduction in Live Pancreatic Islets. Biophysical Journal 111(2): 409–17, Doi: 10.1016/j.bpj.2016.06.014.

[21] Elliott, A.D., Ustione, A., Piston, D.W., 2015. Somatostatin and insulin mediate glucose-inhibited glucagon secretion in the pancreatic α-cell by lowering cAMP. American Journal of Physiology-Endocrinology and Metabolism 308(2): E130–43, Doi: 10.1152/ajpendo.00344.2014.

[22] De Marinis, Y.Z., Salehi, A., Ward, C.E., Zhang, Q., Abdulkader, F., Bengtsson, M., et al., 2010. GLP-1 Inhibits and Adrenaline Stimulates Glucagon Release by Differential Modulation of N-and L-Type Ca2+ Channel-Dependent Exocytosis. Cell Metabolism 11(6): 543–53, Doi: 10.1016/j.cmet.2010.04.007.

[23] Tengholm, A., Gylfe, E., 2017. cAMP signalling in insulin and glucagon secretion: TENGHOLM AND GYLFE. Diabetes, Obesity and Metabolism 19: 42–53, Doi: 10.1111/dom.12993.

[24] Wendt, A., Birnir, B., Buschard, K., Gromada, J., Salehi, A., Sewing, S., et al., 2004. Glucose Inhibition of Glucagon Secretion From Rat α-Cells Is Mediated by GABA Released From Neighboring β-Cells. Diabetes 53(4): 1038–45, Doi: 10.2337/diabetes.53.4.1038.

[25] Ishihara, H., Maechler, P., Gjinovci, A., Herrera, P.-L., Wollheim, C.B., 2003. Islet β-cell secretion determines glucagon release from neighbouring α-cells. Nature Cell Biology 5(4): 330–5, Doi: 10.1038/ncb951.

[26] Rorsman, P., Berggren, P.-O., Bokvist, K., Ericson, H., Möhler, H., Östenson, C.-G., et al., 1989. Glucose-inhibition of glucagon secretion involves activation of GABAA-receptor chloride channels. Nature 341(6239): 233–6, Doi: 10.1038/341233a0.

[27] Almaça, J., Molina, J., Menegaz, D., Pronin, A.N., Tamayo, A., Slepak, V., et al., 2016. Human Beta Cells Produce and Release Serotonin to Inhibit Glucagon Secretion from Alpha Cells. Cell Reports 17(12): 3281–91, Doi: 10.1016/j.celrep.2016.11.072.

[28] Yu, Q., Shuai, H., Ahooghalandari, P., Gylfe, E., Tengholm, A., 2019. Glucose controls glucagon secretion by directly modulating cAMP in alpha cells. Diabetologia 62(7): 1212–24, Doi: 10.1007/s00125-019-4857-6.

[29] Vergari, E., Knudsen, J.G., Ramracheya, R., Salehi, A., Zhang, Q., Adam, J., et al., 2019. Insulin inhibits glucagon release by SGLT2-induced stimulation of somatostatin secretion. Nature Communications 10(1): 139, Doi: 10.1038/s41467-018-08193-8.

[30] Rorsman, P., Huising, M.O., 2018. The somatostatin-secreting pancreatic δ-cell in health and disease. Nature Reviews Endocrinology 14(7): 404–14, Doi: 10.1038/s41574-018-0020-6.

[31] van der Meulen, T., Donaldson, C.J., Cáceres, E., Hunter, A.E., Cowing-Zitron, C., Pound, L.D., et al., 2015. Urocortin3 mediates somatostatin-dependent negative feedback control of insulin secretion. Nature Medicine 21(7): 769–76, Doi: 10.1038/nm.3872.

[32] Kailey, B., van de Bunt, M., Cheley, S., Johnson, P.R., MacDonald, P.E., Gloyn, A.L., et al., 2012. SSTR2 is the functionally dominant somatostatin receptor in human pancreatic β- and α-cells. American Journal of Physiology-Endocrinology and Metabolism 303(9): E1107–16, Doi: 10.1152/ajpendo.00207.2012.

[33] Vieira, E., Salehi, A., Gylfe, E., 2007. Glucose inhibits glucagon secretion by a direct effect on mouse pancreatic alpha cells. Diabetologia 50(2): 370–9, Doi: 10.1007/s00125-006-0511-1.

[34] Cheng-Xue, R., Gómez-Ruiz, A., Antoine, N., Noël, L.A., Chae, H.-Y., Ravier, M.A., et al., 2013. Tolbutamide Controls Glucagon Release From Mouse Islets Differently Than Glucose. Diabetes 62(5): 1612–22, Doi: 10.2337/db12-0347.

[35] Bahl, V., Lee May, C., Perez, A., Glaser, B., Kaestner, K.H., 2021. Genetic activation of α-cell glucokinase in mice causes enhanced glucose-suppression of glucagon secretion during normal and diabetic states. Molecular Metabolism 49: 101193, Doi: 10.1016/j.molmet.2021.101193.

[36] Moede, T., Leibiger, B., Vaca Sanchez, P., Daré, E., Köhler, M., Muhandiramlage, T.P., et al., 2020. Glucokinase intrinsically regulates glucose sensing and glucagon secretion in pancreatic alpha cells. Scientific Reports 10(1): 20145, Doi: 10.1038/s41598-020-76863-z.

[37] Basco, D., Zhang, Q., Salehi, A., Tarasov, A., Dolci, W., Herrera, P., et al., 2018. α-cell glucokinase suppresses glucose-regulated glucagon secretion. Nature Communications 9(1): 546, Doi: 10.1038/s41467-018-03034-0.

[38] Ho, T., Potapenko, E., Davis, D.B., Merrins, M.J., 2023. A plasma membrane-associated glycolytic metabolon is functionally coupled to KATP channels in pancreatic α and β cells from humans and mice. Cell Reports 42(4): 112394, Doi: 10.1016/j.celrep.2023.112394.

[39] Dai, X.-Q., Camunas-Soler, J., Briant, L.J.B., dos Santos, T., Spigelman, A.F., Walker, E.M., et al., 2022. Heterogenous impairment of α cell function in type 2 diabetes is linked to cell maturation state. Cell Metabolism 34(2): 256–268.e5, Doi: 10.1016/j.cmet.2021.12.021.

[40] Briant, L.J.B., Dodd, M.S., Chibalina, M.V., Rorsman, N.J.G., Johnson, P.R.V., Carmeliet, P., et al., 2018. CPT1a-Dependent Long-Chain Fatty Acid Oxidation Contributes to Maintaining Glucagon Secretion from Pancreatic Islets. Cell Reports 23(11): 3300–11, Doi: 10.1016/j.celrep.2018.05.035.

[41] Armour, S.L., Frueh, A., Chibalina, M.V., Dou, H., Argemi-Muntadas, L., Hamiltion, A., et al., 2023. Glucose controls glucagon secretion by regulating fatty acid oxidation in pancreatic alpha cells. Diabetes: db230056, Doi: 10.2337/db23-0056.

[42] Rocha, D.M., Faloona, G.R., Unger, R.H., 1972. Glucagon-stimulating activity of 20 amino acids in dogs. Journal of Clinical Investigation 51(9): 2346–51, Doi: 10.1172/JCI107046.

[43] Gerich, J.E., Frankel, B.J., Fanska, R., West, L., Forsham, P.H., Grodsky, G.M., 1974. Calcium Dependency of Glucagon Secretion from the in Vitro Perfused Rat Pancreas. Endocrinology 94(5): 1381–5, Doi: 10.1210/endo-94-5-1381.

[44] Dean, E.D., Li, M., Prasad, N., Wisniewski, S.N., Von Deylen, A., Spaeth, J., et al., 2017. Interrupted Glucagon Signaling Reveals Hepatic α Cell Axis and Role for L-Glutamine in α Cell Proliferation. Cell Metabolism 25(6): 1362–1373.e5, Doi: 10.1016/j.cmet.2017.05.011.

[45] Gameiro, A., Reimann, F., Habib, A.M., O’Malley, D., Williams, L., Simpson, A.K., et al., 2005. The neurotransmitters glycine and GABA stimulate glucagon-like peptide-1 release from the GLUTag cell line: GLP-1 release triggered by glycine and GABA. The Journal of Physiology 569(3): 761–72, Doi: 10.1113/jphysiol.2005.098962.

[46] Cabrera, O., Jacques-Silva, M.C., Speier, S., Yang, S.-N., Köhler, M., Fachado, A., et al., 2008. Glutamate Is a Positive Autocrine Signal for Glucagon Release. Cell Metabolism 7(6): 545–54, Doi: 10.1016/j.cmet.2008.03.004.

[47] Stanley, C.A., Lieu, Y.K., Hsu, B.Y.L., Burlina, A.B., Greenberg, C.R., Hopwood, N.J., et al., 1998. Hyperinsulinism and Hyperammonemia in Infants with Regulatory Mutations of the Glutamate Dehydrogenase Gene. New England Journal of Medicine 338(19): 1352–7, Doi: 10.1056/NEJM199805073381904.

[48] Lai, B.-K., Chae, H., Gómez-Ruiz, A., Cheng, P., Gallo, P., Antoine, N., et al., 2018. Somatostatin Is Only Partly Required for the Glucagonostatic Effect of Glucose but Is Necessary for the Glucagonostatic Effect of K ATP Channel Blockers. Diabetes 67(11): 2239–53, Doi: 10.2337/db17-0880.

[49] DiGruccio, M.R., Mawla, A.M., Donaldson, C.J., Noguchi, G.M., Vaughan, J., Cowing-Zitron, C., et al., 2016. Comprehensive alpha, beta and delta cell transcriptomes reveal that ghrelin selectively activates delta cells and promotes somatostatin release from pancreatic islets. Molecular Metabolism 5(7): 449–58, Doi: 10.1016/j.molmet.2016.04.007.

[50] Trube, G., Rorsman, P., Ohno-Shosaku, T., 1986. Opposite effects of tolbutamide and diazoxide on the ATP-dependent K+ channel in mouse pancreatic beta-cells. Pfligers Archiv European Journal of Physiology 407(5): 493–9, Doi: 10.1007/BF00657506.

[51] Braun, M., Ramracheya, R., Amisten, S., Bengtsson, M., Moritoh, Y., Zhang, Q., et al., 2009. Somatostatin release, electrical activity, membrane currents and exocytosis in human pancreatic delta cells. Diabetologia 52(8): 1566–78, Doi: 10.1007/s00125-009-1382-z.

[52] Schuit, F., De Vos, A., Farfari, S., Moens, K., Pipeleers, D., Brun, T., et al., 1997. Metabolic fate of glucose in purified islet cells. Glucose-regulated anaplerosis in beta cells. The Journal of Biological Chemistry 272(30): 18572–9, Doi: 10.1074/jbc.272.30.18572.

[53] Li, C., Najafi, H., Daikhin, Y., Nissim, I.B., Collins, H.W., Yudkoff, M., et al., 2003. Regulation of leucine-stimulated insulin secretion and glutamine metabolism in isolated rat islets. The Journal of Biological Chemistry 278(5): 2853–8, Doi: 10.1074/jbc.M210577200.

[54] Zhou, Y., Jetton, T.L., Goshorn, S., Lynch, C.J., She, P., 2010. Transamination is required for {alpha}-ketoisocaproate but not leucine to stimulate insulin secretion. The Journal of Biological Chemistry 285(44): 33718–26, Doi: 10.1074/jbc.M110.136846.

[55] Gao, Z., Young, R.A., Li, G., Najafi, H., Buettger, C., Sukumvanich, S.S., et al., 2003. Distinguishing features of leucine and alpha-ketoisocaproate sensing in pancreatic beta-cells. Endocrinology 144(5): 1949–57, Doi: 10.1210/en.2002-0072.

[56] Svendsen, B., Larsen, O., Gabe, M.B.N., Christiansen, C.B., Rosenkilde, M.M., Drucker, D.J., et al., 2018. Insulin Secretion Depends on Intra-islet Glucagon Signaling. Cell Reports 25(5): 1127–1134.e2, Doi: 10.1016/j.celrep.2018.10.018.

[57] Rodriguez-Diaz, R., Tamayo, A., Hara, M., Caicedo, A., 2020. The Local Paracrine Actions of the Pancreatic α-Cell. Diabetes 69(4): 550–8, Doi: 10.2337/dbi19-0002.

[58] Holter, M.M., Saikia, M., Cummings, B.P., 2022. Alpha-cell paracrine signaling in the regulation of beta-cell insulin secretion. Frontiers in Endocrinology 13: 934775, Doi: 10.3389/fendo.2022.934775.

[59] Campbell, J.E., Newgard, C.B., 2021. Mechanisms controlling pancreatic islet cell function in insulin secretion. Nature Reviews Molecular Cell Biology 22(2): 142–58, Doi: 10.1038/s41580-020-00317-7.

[60] Huising, M.O., 2020. Paracrine regulation of insulin secretion. Diabetologia 63(10): 2057–63, Doi: 10.1007/s00125-020-05213-5.

[61] MacDonald, M.J., 2002. Differences between mouse and rat pancreatic islets: succinate responsiveness, malic enzyme, and anaplerosis. American Journal of Physiology. Endocrinology and Metabolism 283(2): E302–310, Doi: 10.1152/ajpendo.00041.2002.

[62] Lewandowski, S.L., Cardone, R.L., Foster, H.R., Ho, T., Potapenko, E., Poudel, C., et al., 2020. Pyruvate Kinase Controls Signal Strength in the Insulin Secretory Pathway. Cell Metabolism 32(5): 736–750.e5, Doi: 10.1016/j.cmet.2020.10.007.

[63] Zaborska, K.E., Dadi, P.K., Dickerson, M.T., Nakhe, A.Y., Thorson, A.S., Schaub, C.M., et al., 2020. Lactate activation of α-cell KATP channels inhibits glucagon secretion by hyperpolarizing the membrane potential and reducing Ca2+ entry. Molecular Metabolism 42: 101056, Doi: 10.1016/j.molmet.2020.101056.

[64] Ravier, M.A., Rutter, G.A., 2005. Glucose or insulin, but not zinc ions, inhibit glucagon secretion from mouse pancreatic alpha-cells. Diabetes 54(6): 1789–97, Doi: 10.2337/diabetes.54.6.1789.

[65] Rutter, G.A., Sidarala, V., Kaufman, B.A., Soleimanpour, S.A., 2023. Mitochondrial metabolism and dynamics in pancreatic beta cell glucose sensing. The Biochemical Journal 480(11): 773–89, Doi: 10.1042/BCJ20230167.

[66] Foster, H.R., Ho, T., Potapenko, E., Sdao, S.M., Huang, S.M., Lewandowski, S.L., et al., 2022. β-cell deletion of the PKm1 and PKm2 isoforms of pyruvate kinase in mice reveals their essential role as nutrient sensors for the KATP channel. ELife 11: e79422, Doi: 10.7554/eLife.79422.

[67] Abulizi, A., Cardone, R.L., Stark, R., Lewandowski, S.L., Zhao, X., Hillion, J., et al., 2020. Multi-Tissue Acceleration of the Mitochondrial Phosphoenolpyruvate Cycle Improves Whole-Body Metabolic Health. Cell Metabolism 32(5): 751–766.e11, Doi: 10.1016/j.cmet.2020.10.006.

[68] Merrins, M.J., Corkey, B.E., Kibbey, R.G., Prentki, M., 2022. Metabolic cycles and signals for insulin secretion. Cell Metabolism: S1550-4131(22)00223–6, Doi: 10.1016/j.cmet.2022.06.003.

[69] Armour, S.L., Stanley, J.E., Cantley, J., Dean, E.D., Knudsen, J.G., 2023. Metabolic regulation of glucagon secretion. The Journal of Endocrinology: JOE-23-0081, Doi: 10.1530/JOE-23-0081.

[70] Riahi, Y., Kogot-Levin, A., Kadosh, L., Agranovich, B., Malka, A., Assa, M., et al., 2023. Hyperglucagonaemia in diabetes: altered amino acid metabolism triggers mTORC1 activation, which drives glucagon production. Diabetologia, Doi: 10.1007/s00125-023-05967-8.

[71] Stanley, C.A., Lieu, Y.K., Hsu, B.Y., Burlina, A.B., Greenberg, C.R., Hopwood, N.J., et al., 1998. Hyperinsulinism and hyperammonemia in infants with regulatory mutations of the glutamate dehydrogenase gene. The New England Journal of Medicine 338(19): 1352–7, Doi: 10.1056/NEJM199805073381904.

[72] Ackermann, A.M., Zhang, J., Heller, A., Briker, A., Kaestner, K.H., 2017. High-fidelity Glucagon-CreER mouse line generated by CRISPR-Cas9 assisted gene targeting. Molecular Metabolism 6(3): 236–44, Doi: 10.1016/j.molmet.2017.01.003.

[73] Ahn, S.-H., Granger, A., Rankin, M.M., Lam, C.J., Cox, A.R., Kushner, J.A., 2019. Tamoxifen suppresses pancreatic β-cell proliferation in mice. PloS One 14(9): e0214829, Doi: 10.1371/journal.pone.0214829.

[74] Carboneau, B.A., Le, T.D.V., Dunn, J.C., Gannon, M., 2016. Unexpected effects of the MIP-CreER transgene and tamoxifen on β-cell growth in C57Bl6/J male mice. Physiological Reports 4(18), Doi: 10.14814/phy2.12863.

[75] Madisen, L., Garner, A.R., Shimaoka, D., Chuong, A.S., Klapoetke, N.C., Li, L., et al., 2015. Transgenic mice for intersectional targeting of neural sensors and effectors with high specificity and performance. Neuron 85(5): 942–58, Doi: 10.1016/j.neuron.2015.02.022.

[76] Muntean, B.S., Zucca, S., MacMullen, C.M., Dao, M.T., Johnston, C., Iwamoto, H., et al., 2018. Interrogating the Spatiotemporal Landscape of Neuromodulatory GPCR Signaling by Real-Time Imaging of cAMP in Intact Neurons and Circuits. Cell Reports 22(1): 255–68, Doi: 10.1016/j.celrep.2017.12.022.

[77] Gregg, T., Poudel, C., Schmidt, B.A., Dhillon, R.S., Sdao, S.M., Truchan, N.A., et al., 2016. Pancreatic β-Cells From Mice Offset Age-Associated Mitochondrial Deficiency With Reduced KATP Channel Activity. Diabetes 65(9): 2700–10, Doi: 10.2337/db16-0432.

