## Supplemental Figure 1, Table 1, and Table 2 for "Leucine suppresses glucagon secretion from pancreatic islets by directly modulating α-cell cAMP"

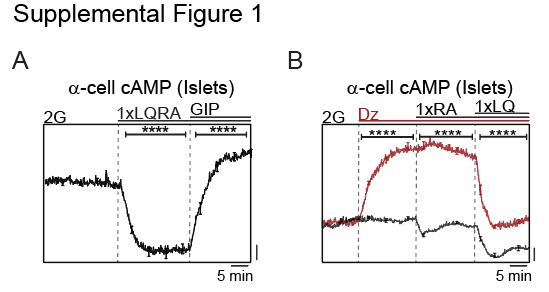

**Supplemental Figure 1.** (A) Epifluorescence microscopy of α-cell cAMP in intact mouse islets stimulated with 2 mM glucose (2G), amino acids provided at physiological concentration (L, 0.5 mM leucine; Q, 0.6 mM glutamine; R, 0.2 mM arginine; A, 2.1 mM alanine), and GIP (10 nM) as indicated. Data reflect 147 islets from n = 4 mice. (B) Epifluorescence microscopy of α-cell cAMP in intact mouse islets stimulated with 2 mM glucose (2G), amino acids provided at physiological concentration (L, 0.5 mM leucine; Q, 0.6 mM glutamine; R, 0.2 mM arginine; A, 2.1 mM alanine), and 200 μM diazoxide (Dz) as indicated. Data reflect 147 islets from n = 4 mice. Data are shown as mean ± SEM for each condition with ****p < 0.0001 by Student’s t test.

**Supplemental Table 1.** Summary characteristics of human islet donors. BMI, body mass index; HbA1c, glycated hemoglobin.

| ***Donor ID*** | ***Age***  ***(years)*** | ***Sex*** | ***BMI*** | ***HbA1c*** |
| --- | --- | --- | --- | --- |
| A480 | 54 | F | 27.5 | 5.2% |
| P301 | 63 | F | 22.1 | 5.6% |
| I313 | 28 | M | 29.3 | 5.0% |
| I325 | 46 | M | 39.2 | 5.6% |

**Supplemental Table 2.** Qualitative determination of reporter expression observed in the islets from GcgCre^ERT^:GCaMP6s mice. Spurious expression of GCaMP6s was observed, *i.e.* in the absence of tamoxifen or 5-hydroxytamoxifen, when GcgCre^ERT^ was present in the dam (n = 10 mice from 4 breeding pairs). No expression of GCaMP6s was observed in the absence of GcgCre^ERT^.

| ***Sire ID*** |  |  | ***Dam ID*** |  |  | ***Mouse ID*** | ***Leaky GCaMP6s?*** |
| --- | --- | --- | --- | --- | --- | --- | --- |
| 373793 | GcgCreERT | -/+ | 397258 | GCaMP6s | -/- | 439632 | No |
| 373793 | GcgCreERT | -/+ | 397258 | GCaMP6s | -/- | 439633 | No |
| 373793 | GcgCreERT | -/+ | 397258 | GCaMP6s | -/- | 446526 | No |
| 373793 | GcgCreERT | -/+ | 397258 | GCaMP6s | -/- | 446527 | No |
| 428931 | GCaMP6s | -/- | 416156 | GcgCreERT | -/+ | 452992 | Yes |
| 428932 | GCaMP6s | -/- | 419479 | GcgCreERT | -/+ | 453000 | Yes |
| 428932 | GCaMP6s | -/- | 419479 | GcgCreERT | -/+ | 453001 | Yes |
| 428932 | GCaMP6s | -/- | 419479 | GcgCreERT | -/+ | 453002 | Yes |
| 428932 | GCaMP6s | -/- | 419479 | GcgCreERT | -/+ | 453003 | Yes |
| 373793 | GcgCreERT | -/+ | 397258 | GCaMP6s | -/- | 454714 | No |
